# Multi-omics framework to reveal the molecular determinants of fermentation performance in wine yeast populations

**DOI:** 10.1101/2023.12.02.569693

**Authors:** Miguel de Celis, Javier Ruiz, Belen Benitez-Dominguez, Javier Vicente, Sandra Tomasi, Sergio Izquierdo-Gea, Nicolás Rozés, Candela Ruiz-de-Vila, Jordi Gombau, Fernando Zamora, Alicia Barroso, Laura C. Terron-Camero, Eduardo Andres-Leon, Antonio Santos, Ignacio Belda

## Abstract

**Background:** Connecting the composition and function of industrial microbiomes is a major aspiration in microbial biotechnology. Here, we address this question in wine fermentation, a model system where the diversity and functioning of fermenting yeast species is determinant of the flavor and quality of the resulting wines.

**Results:** First, we surveyed yeast communities associated with grape musts collected across wine appellations, revealing the importance of environmental (i.e., biogeography) and anthropic factors (i.e., farming system) in shaping community composition and structure. Then, we assayed the fermenting yeast communities in synthetic grape must under common winemaking conditions. The dominating yeast species defines the fermentation performance and metabolite profile of the resulting wines, and it is determined by the initial fungal community composition rather than the imposed fermentation conditions. Yeast dominance also had a more pronounced impact on wine meta-transcriptome than fermentation conditions. We unveiled yeast-specific transcriptomic profiles, leveraging different molecular functioning strategies in wine fermentation environments. We further studied the orthologs responsible for metabolite production, revealing modules associated with the dominance of specific yeast species. This emphasizes the unique contributions of yeast species to wine flavor, here summarized in an array of orthologs that defines the individual contribution of yeast species to wine ecosystem functioning.

**Conclusions:** Our study bridges the gap between yeast community composition and wine metabolite production, providing insights to harness diverse yeast functionalities with the final aim to producing tailored high-quality wines.

## Background

Originally, wine fermentations were spontaneous and the native yeast communities present in the grape surface were responsible for completing the fermentation of grape musts. The grape surface constitutes a complex microbial ecosystem with yeasts, filamentous fungi and bacteria, each influencing wine production differently [1]. Yeast communities play crucial roles in determining the chemical and sensory properties of the resulting wines [2–4]. A diverse yeast microbiome in grape musts can enhance the aromatic complexity of final wines [5–8]. The inherent diversity in grape musts, coupled with the spontaneous nature of fermentations, implies limited control over population dynamics during wine fermentation, influencing its kinetics and final sensory output. In contrast to the spontaneity of natural fermentations, the wine industry introduced the standardization of wine fermentations through the inoculation of commercial yeast strains or consortia with predefined traits [9]. Regardless of the fermentation strategy followed, deciphering the molecular determinants of the individual contribution of wine yeast species to wine flavor is essential to advance in the targeted improvement of wine quality.

Several factors influence yeast communities associated with grape berries, subsequently shaping the diversity of fermenting yeast species in grape musts. Edaphoclimatic factors and vineyard management practices contribute to the complexity of these communities [10,11]. After crushing grapes, these microbial populations have to cope with various environmental challenges during the fermentation of grape musts, such as high osmotic pressure, low pH, suboptimal temperatures for growth, increasing ethanol concentrations, and anaerobic conditions [12,13]. This leads to a rapid succession of yeast populations where the initially wide fungal diversity is wiped out by the ethanol toxicity and is replaced by ethanol-tolerant fermentative yeasts, mainly *Saccharomyces cerevisiae* [14,15]. By changing temperature, or adding sulfur dioxide (SO_2_) or nitrogen nutrients, winemakers have the possibility to modify the fermentation dynamics and affecting the yeast performance [16–18]. Understanding the influence of these conditions on yeast growth and fermentation processes will enhance future achievements in the oenological industry, facilitating a more rational use of yeast cultures, food additives, and nutrients in wine fermentations. However, to gain control and predict the metabolite profile of final wines, it is necessary to initially comprehend the functional potential of wild yeast communities subjected to varying fermentation conditions, considering common industrial practices and the diverse environmental conditions of grapevines and fermenting facilities influencing winemaking process.

In this study, we combined observational studies with laboratory fermentations to assess factors shaping yeast community composition associated with grape berries and to identify functional yeasts associated with wine metabolite production under different fermentation conditions. We integrated multi-omics data to understand the connection between the composition and function of fermenting yeast communities with the final metabolite profile of wines. In doing so, we aim to contribute to our ecological understanding of wine ecosystems and highlight the importance of preserving the complex dynamics even in controlled fermentation processes. To do so, we surveyed five distinct Spanish wine appellations, sampling grapes from vineyards under conventional and organic management. We also sampled at lower geographic scales to finely disentangle whether grape must and fungal variability was distance dependent. Specifically, for one region (La Rioja) we surveyed 3 conventional and 3 organic vineyards, one of each sampled per triplicate, in three different vine rows (**Supplementary Figure S1**). We analyzed the metabolite and fungal community profiles of fresh grape musts, defining their initial variability, and subject them to spontaneous fermentations in four different fermentation conditions. Then, we inoculated the fermenting yeast communities in synthetic grape must, and studied the associations between their functional profiles, through meta-transcriptomic analysis, and the final metabolite composition of the resulting wines.

## Methods

In this study, we carried an observational study aiming to assess the variability, and the driving factors, of grape must and its associated yeast communities across different conventionally and organic managed vineyards, sampling across regional and local locations to determine if it is distance dependent. Then, with the variability of yeast communities sampled, we carried laboratory fermentations and performed multi-omics analysis to understand the connection between the composition and function of fermenting yeast communities with the final metabolite profile of wines.

### Observational study design

A total of nine different locations were sampled from five different Spanish wine appellations (**Supplementary Figure S1**). The wine appellations were Ribera del Guadiana (RdG), Valdepeñas (VLP), La Mancha (LM), Madrid (M), and La Rioja (R). Within La Rioja we sampled three locations (R1, R2 and R3; the later per triplicate, in three different rows within the same plot: R3A, R3B, R3C). This spatial survey was applied in parallel, sampling two vineyards within each location, under conventional and under organic farming, respectively. Contrary to conventional management, organic vineyards are restrictive with the use of inorganic fertilizers as well as phytosanitary products. The regulatory specifications outlining the criteria for Conventional and Organic classifications are established according to the European Union Regulation (EC) No. 1234/2007 (Official Journal of the European Union, 2007). The different patterns of fungal community and metabolite production during fermentation are largely dependent on grape variety [10]. Therefore, we only sampled grapes of Tempranillo variety, when possible, that is, in all cases except in conventional farming in RdG, where we could only sample grapes of Red Garnacha variety. Since the physical-chemical composition of Red Garnacha grapes was greatly different than Tempranillo grapes (**Supplementary Figure S2**) we only used RdG-conventional sample for fungal community isolation.

### Grape processing and grape must fermentations

We collected 3 kg of grapes (*Vitis vinifera* L. cv. Tempranillo, except RdG-conventional sample which was Red Garnacha variety) in 5 bunches of grapes from five different grapevine plants, making a composite sample from each sampling point. Upon arrival at the laboratory, grapes were pressed under sterile conditions and the resulting grape must was dispensed into four sterile glass bottles. Immediately after filling the bottles, an initial sample was collected for DNA extraction and sequencing to assess the initial fungal community composition. Then, these bottles were subjected to fermentation under different conditions: i) Control condition; ii) 18°C condition, fermented at lower temperature (18°C compared to the 25°C of the control condition); iii) NH_4_ condition, supplemented with 300 mg/L diammonium sulfate ((NH_4_)_2_SO_4_); and iv) SO_2_ condition, adding 100 mg/L of potassium metabisulfite (K_2_S_2_O_5_). We defined the end of wine fermentations as the point when the weight loss remained consistently below 0.01 g per day for two consecutive days. At this point collected samples for DNA extraction and sequencing to assess the final fungal community composition.

### Laboratory fermentations

To finely assess the impact of fermentation conditions in the transcriptional and metabolite patterns of wine yeasts, we replicated the fermentations using synthetic grape must (SGM), prepared as described by Ruiz et al., [19]. To obtain the greatest diversity of wild communities to further inoculate the experimental fermentations in SGM, we extracted 2 mL of fermenting grape musts at the tumultuous stage (between 23 and 45% of total sugars consumed, **Supplementary Table S1**) from each control replicate of the 18 fermentation assays performed in the natural grape musts (**Supplementary Figure S1**). These 18 samples were frozen (−80°C) and then, after thawing, centrifugation, resuspension, and standardization of the Optical Density (OD_600mm_) of the resulting pellet, they were used as seed communities to inoculate the SGM. Then, we inoculated the seed communities in 80 mL of SGM bottles, per quadruplicate. These bottles were subjected to different fermentative conditions (**Supplementary Figure S1**): i) Control condition; ii) 18°C condition, fermented at lower temperature (18°C compared to the 25°C of the control condition); iii) NH_4_ condition, supplemented with 300 mg/L diammonium sulfate ((NH_4_)_2_SO_4_); and iv) SO_2_ condition, adding 100 mg/L of potassium metabisulfite (K_2_S_2_O_5_). As with spontaneous fermentations, we defined the end of wine fermentations as the point when the weight loss remained consistently below 0.01 g per day for two consecutive days.

### Fungal community assessment

#### Amplicon sequencing

We used the DNeasy PowerSoil Pro Kit (Qiagen) for DNA extraction following manufacturer’s instructions. We checked DNA quality and quantity using a NanoDrop 2000 (Thermo Fisher Scientific, USA) and Qubit Fluorometer (Thermo Fisher Scientific, USA), respectively. Then, the diversity and composition of the fungal community were determined by amplicon sequencing. DNA sequencing was performed at the “López-Neyra” Institute of Parasitology and Biomedicine. For library preparation, we used ITS2_fITS7 forward (TCCTCCGCTTATTGATATGC) and ITS4 reverse (GTGARTCATCGAATCTTTG) primers. Libraries were subsequently sequenced on Illumina® MiSeq instrument using 2 × 300 paired-end reads as per the manufacturer’s instructions.

We obtained a total of 17098692 good quality sequences, averaging 62863 ± 20796 per sample (further quality analysis at https://github.com/Migueldc1/Wineteractions/). Sequence analysis was performed with *dada2* v1.24.0 [20] R package, after FastQC [21] and MultiQC quality assessment [22]. Dada2 identifies amplicon sequence variants (ASVs), allowing us to distinguish true biological variation from sequencing errors and PCR artefacts, and changes of one nucleotide can be detected [20]. This allows for more accurate and precise identification of unique microbial taxa and quantification of their abundances. Primers were removed using *cutadapt* [23], and *Biostrings* [24] R package to assess their orientation in the reads. After quality assessment, low-quality ends were deleted, and no mismatches were allowed when merging paired reads. Once chimaeras were removed, we finally assigned taxonomy to our ITS reads using the UNITE v9.0 database [25].

#### Diversity and statistical analysis

We first addressed whether there were differences in alpha diversity among the different geographical origins and between farming practices. To do so we used Hill-based diversity index in which the importance given to the relative abundance of each ASV can be varied [26]. This importance is determined by the diversity order (*q*). For instance, when *q* = 0, the relative abundance is not considered, and hence, the value equals the richness. When *q* = 1, each ASV is weighted according to their relative abundance, and when *q* = 2 more weight is given to abundant ASVs. These last two values equal the exponential Shannon index (Hill-Shannon), and reciprocal Simpson index (Hill-Simpson) [27]. We calculated the Hill-based alpha diversity using the *hillR* R package [28]. To disentangle differences in alpha diversity caused by different farming practices and geographical origin, we used linear mixed models with the origin as a random factor and the farming practices as fixed factors, using the *nlme* R package [29].

Differences in community composition across the studied geographical origin and farming practices (β-diversity) were evaluated with the *vegan* R package [30]. We first calculated Bray-Curtis dissimilarity matrices from the ASV table collapsed at the genus level, to consider relative abundance of genera in our samples. Then, we performed multivariate permutational multivariate analysis of variance (PERMANOVA) to assess the effect of origin and farming on community composition, and non-metric multidimensional scaling (NMDS) to compress dimensionality into two dimensional plots. Code for statistical analyses is available at https://github.com/Migueldc1/Wineteractions.

### Metabolite profiling

For initial grape must samples we measured pH, and concentration of non-volatile compounds: organic (amino acid related) and inorganic (ammonium) nitrogen, sugar (glucose and fructose), and L-malic acid. Due to the increased metabolite complexity of wines [31], we analyzed non-volatile compounds also including ethanol, acetic acid, L-lactic acid, tartaric acid, citric acid, succinic acid, and glycerol; and volatile compounds: ethyl acetate, fusel alcohols (isopropanol, 1-propanol, 2-methyl-propanol, 1-butanol, 2-methyl-1-butanol, 1-hexanol, 2-ethyl-1-hexanol, 2-butanol and 2-phenylethanol), fusel alcohol acetates (isobutyl acetate, isoamyl acetate, hexyl acetate and 2-phenylethanol acetate), ethyl-esters of fatty acids (EEFA: ethyl butanoate, ethyl octanoate and ethyl dodecanoate), short-chain fatty acids (SCFA: propionic acid, isobutyric acid, butyric acid, valeric acid and 2-methylbutanoic acid), and medium-chain fatty acids (MCFA: hexanoic acid, octanoic acid and decanoic acid) of fermented SGM samples. The pH was measured using a pH meter (Crison pH Meter Basic 20, Crison-Spain) and the concentrations of non-volatile compounds were measured using specialized enzymatic kits and the analyzer Y15 (Biosystems, Spain) following manufacturer’s instructions. Gas chromatography-mass spectrometry (GC-FID) was used to measure the concentration of volatile compounds as previously described [32]. Raw data detailed in **Additional File 2**.

### Metatranscriptomic analysis

#### RNA extraction and sequencing

We collected SGM samples during the tumultuous stage of fermentation (here with between 5 and 54% of sugars consumed, **Supplementary Table S1**) and centrifuged at 7,000 rpm and 4°C for 5 min. Biomass was quickly frozen with liquid nitrogen and stored at −80°C until RNA extraction. RNA extraction protocol was carried out according to the specifications provided in the Quick-RNA Fungal/Bacterial MicroPrep kit (ZymoResearch). RNA quality analysis, library preparation, RNA sequencing and bioinformatics analyses were carried out at the Bioinformatics and Genomics Unit of The López-Neyra Institute of Parasitology and Biomedicine (IPBLN-CSIC, Granada, Spain). The quality of the RNAs was evaluated using Bioanalyzer (Agilent Technologies) and samples with RNA Integrity Number (RIN) ≥ 8.2 were selected for further analysis (https://github.com/Migueldc1/Wineteractions/Quality_Assessment). Next, rRNA was depleted using the NEBNext rRNA Depletion Kit (Human/Mouse/Rat; New England Biolab) following manufacturer’s specifications. Finally, libraries were constructed using TruSeq^TM^ Stranded RNA sample preparation kit, according to Illumina’s instructions. In addition, libraries quality was validated by Qubit dsDNA HS Assay Kit (ThermoFisher) and 2100 Bioanalyzer (Agilent Technologies). Afterwards, these libraries were sequenced on an Illumina NextSeq High Output, producing 75 bp paired end reads. We obtained a total of 15720876 ± 1906845 for further bioinformatic analysis (quality assessment can be found at https://github.com/Migueldc1/Wineteractions/Quality_Assessment).

#### Bioinformatic analysis

The quality assessment of the raw reads from RNA-sequencing was performed with FastQC [21] and MultiQC software [22]. Overrepresented rRNA fragments were removed using sortmeRNA software [33]. The clean meta-transcriptomics sequence reads were used to assess the taxonomic composition of the SGM fermentations using Kraken 2 [34], and Bracken [35] and a custom database built with the fungal genomes deposited in RefSeq and *Hanseniaspora* genomes from GenBank (including *H. guilliermondii*, *H. opuntiae*, *H. osmophila*, *H. uvarum*, and *H. vineae*). Besides, non-rRNA reads were assembled into contigs/transcripts with Trans-ABySS v2.0.1 [36] using 21, 29, 39, and 59 k-mers. We then, quality checked the assemblies with Assembly-stats v1.0.1 and BUSCO [37], using the *fungi_odb10* database, to assess how well represented is the functional composition of the active fungal community. Functional annotation of transcripts was carried using the eggNOG-mapper v2.1.3 [38] with the DIAMOND aligner [39] against the eggnog database obtaining ortholog annotation. We used the Burrows-Wheeler aligner with BWA software to align the reads to the transcripts [40], as we are not performing the alignment against a reference genome. Finally, we used FeatureCounts v2.0.1 [41] to obtain the number of read counts per ortholog. Differently Expressed (DE) orthologs among dominant yeasts were calculated accounting for different fermentative conditions and origins, using DESeq2 package v1.26.0 [42]. Here, we considered a yeast species as dominant when its relative abundance exceeded 90%, and for further differential expression analysis we focused on samples dominated by *Hanseniaspora*, *Lachancea* and *Saccharomyces*. An ortholog is considered DE when the False Discovery Rate (FDR) value is < 0.05 and the absolute log2 fold change > 1. After that, we used the GOseq v1.48.0 R package [43] to perform gene ontology enrichment analysis on the DE orthologs. Specifically, it focuses on the biological process category of GO terms. The resulting enriched GO terms were filtered to include only those with FDR < 0.05. Code for bioinformatics and statistical analyses is available at https://github.com/Migueldc1/Wineteractions.

We found discrepancies between taxonomic composition assessed via ITS amplicon sequencing and RNA-sequencing taxonomic assignation (**Supplementary Figure S3**), especially concerning the detection of *Hanseniaspora* through ITS sequencing. DNA-based amplicon methods suffer from important biases such as preferential primer binding (leading to bias for or against certain taxa) or not differentiating between living or dead cells [44,45]. RNA sequencing (RNA-seq) captures actively expressed genes, revealing the relative abundance of active members of the community, and thus, we considered this method for further analysis.

## Results and discussion

### Composition and structure of fungal communities in fresh grape musts

First, we conducted an observational study to identify the variability of yeast communities associated with grape berries across wine appellations, identifying the main factors influencing their composition. Climate is considered a pivotal determinant, influencing grape maturation and quality, and consequently, the associated yeast communities [46,47]. Additionally, viticultural practices, such as conventional or organic farming managements, play crucial roles, with conventional farming supporting higher grape berry yields due to nutrient inputs and phytosanitary products [48]. Indeed, we observed that sample location, encompassing the environmental differences among localities, was the main factor defining the metabolite composition (**Figure 1A**) and fungal diversity (**Figure 1B**) of grape musts (**Supplementary Table S2**, **Supplementary Table S3**). We also observed differences in grape must physical-chemical and fungal composition between conventionally and organically managed samples (**Supplementary Table S2**, **Supplementary Table S3**). This effect of farming regime is especially relevant at the local scale, as observed in samples collected within La Rioja appellation (**Supplementary Table S2**, **Supplementary Table S3**), in agreement with previous findings [49]. These findings underscore the complexity of the grapevine ecosystem and the need for a nuanced understanding of both regional and local factors influencing yeast communities.

**Figure 1.**
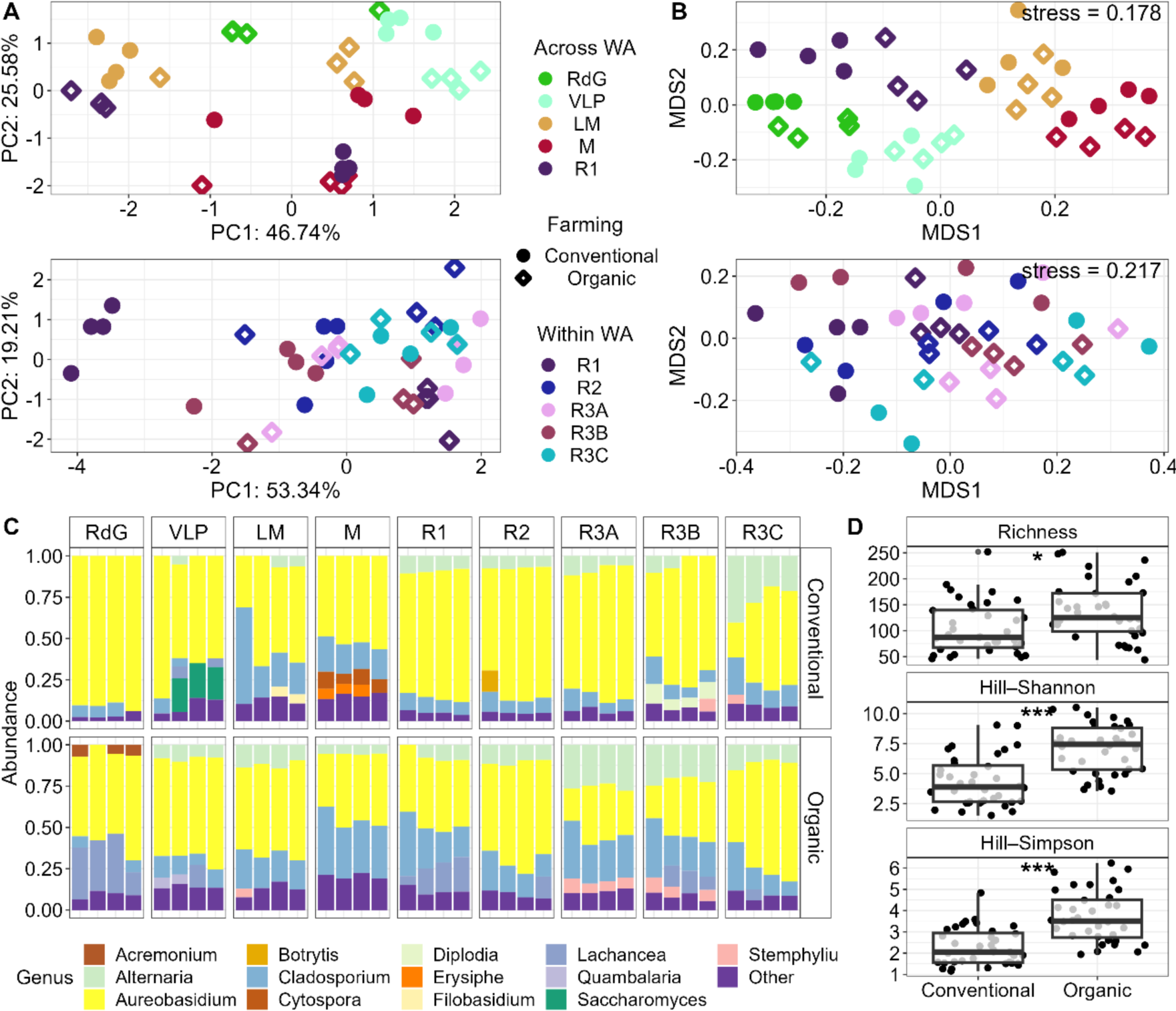
Grape must composition and fungal community description of initial samples. **A)** Principal Component Analysis (PCA) representing grape must composition diversity across (upper panel, n = 4) and within (bottom panel, n = 4) wine appellations (WA). Raw data detailed in **Additional File 2**. **B)** Beta diversity of fungal communities based on a non-metric multidimensional scaling analysis (NMDS, stress parameter indicated in the plot) from Bray-Curtis dissimilarity matrix calculated from genus abundance table. Colors represent the origin of the samples, whereas shapes represent farming practices. **C)** Relative abundance of fungal genus shown for each replicate from each grape sample. “Other” include genus with relative abundance <5%. **D)** Alpha diversity comparison between farming practices (n = 36). The analysis was performed using Hill-based indices with different diversity orders (q). We selected Richness (q = 0), Hill-Shannon (q = 1) and Hill-Simpson (q = 2) to account for the complexity of such communities. We performed Wilcoxon test to evaluate the differences between organic and conventional practices, asterisks denote the significance levels (\*\*\**p* < 0.001, \*\**p* < 0.01 and \**p* < 0.05).

Concerning community composition, the genus *Aureobasidium* dominated all communities, often accompanied by *Cladosporium* and *Lachancea* (**Figure 1C**), consistent with their prevalence in vineyard ecosystems [15,50]. We also identified a high prevalence of *Alternaria,* ubiquitous and predominant filamentous fungi in vineyards worldwide [51]. Samples from vineyards under organic management presented higher fungal diversity, considering phylotype richness and dominant phylotypes (**Figure 1D**). This finding aligns with previous studies attributing the reduced diversity of grape-associated microorganisms in conventional managed vineyards to the adverse impact of phytosanitary products [49]. In this context, *Aureobasidium* could present higher resistance to these products and greater propensity to dominate the fungal communities of grapes from conventional farming regimes, as evidenced by its higher abundance in conventional vineyards (Wilcoxon, *p* < 0.001). The relative abundance of *Cladosporium* and *Lachancea*, on their part, increased in organic vineyards (Wilcoxon, *p* < 0.001). As previously described, *Saccharomyces* is rarely found in fresh grape musts [52]; in our case, it was only detected at relatively high abundances in conventional vineyards from Valdepeñas. Fungal communities associated with grape berries are mainly comprised by filamentous (*Alternaria* or *Cladosporium*) and yeast-like fungus (*Aureobasidium*), ethanol-tolerant yeasts like *Lachancea* or *Saccharomyces* flourish during fermentation [15]. Understanding the diversity of fungal species associated with grape berries is of great interest to infer the range of expectable fermenting communities across wine appellations.

### Manipulating the fermentation environment had no effect on yeast population dynamics

We subjected the obtained fresh grape musts to spontaneous fermentation under four different conditions widely used in wine production (control condition, low temperature, high doses of nitrogen and sulfite addition; see methods for further details). These contrasting fermentation conditions had no major effect on the yeast dynamics during the process, since we found no distinct trend in the dominance of yeast species across conditions (**Supplementary Figure S4A**). *Kluyveromyces*, *Lachancea* and *Saccharomyces* dominated most fermentations, with the initial community variability within sampling location seemingly determining which yeast dominated the fermenting communities. The imposed fermentation conditions also had a minor effect on the metabolite profile of the resulting wines (**Supplementary Figure S4B, Supplementary Table S4**), which were mainly determined by the sample location (which includes the different metabolite composition of the initial grape musts and the dominant yeast population). This kind of studies reignite the debate about the actual possibilities of manipulating spontaneous fermentations by enological interventions, and bring consideration of whether the exclusive avenue for crafting customizable wines lies in the meticulous bottom-up design of synthetic microbial consortia (Ruiz et al., 2023). To better assess the associations between the composition and function of fermenting yeast communities we should move to a fully controlled experimental set up. This would allow us to remove the masking effect of the different physical-chemical environment provided using different fresh grape musts. Thus, we performed parallel laboratory fermentations, using the fermenting yeast communities obtained at the tumultuous stage of the control natural fermentations as seed communities to inoculate synthetic grape musts (**Supplementary Figure S1**).

### Metabolite production by different fermentative yeasts

Our experimental fermentations (Control, 18°C, NH_4_ and SO_2_) were dominated by a handful of yeast genera frequently detected in wine yeast communities [15,53], highlighting *Hanseniaspora*, *Lachancea* and *Saccharomyces* (**Figure 2A**). Performing our assays without the typical contamination of winery facilities, where *Saccharomyces* is part of the resident microbiota, allowed us to observe fermentations dominated by non-*Saccharomyces* yeasts. Fermentations dominated by *Hanseniaspora* and *Lachancea* presented 47.37 ± 16.98% and 16.65 ± 6.54% of unconsumed sugars, respectively (**Figure 2B, Additional File 1**), due to their limited fermentative capacity and ethanol tolerance, preventing them to complete the fermentation process [54–56]. In addition, approximately half of SO_2_ treatments did not even start the fermentation (data not shown), mostly in *Lachancea* or *Hanseniaspora* dominated samples. Different yeast species exhibit varying sensitivity to this antimicrobial, commonly used in wines, with *Saccharomyces* usually displaying higher tolerances [57]. We did not observe a conserved metabolite pattern associated to the fermentation conditions imposed across samples (**Supplementary Figure S5A**, PERMANOVA: R^2^ = 0.062, *p* = 0.421). We only found a general decrease in the final concentration of tartaric and succinic acids when adding diammonium sulfate as a nitrogen nutrient, NH_4_ condition (**Supplementary Figure S6**, further discussion can be found in **Additional File 2**).

**Figure 2.**
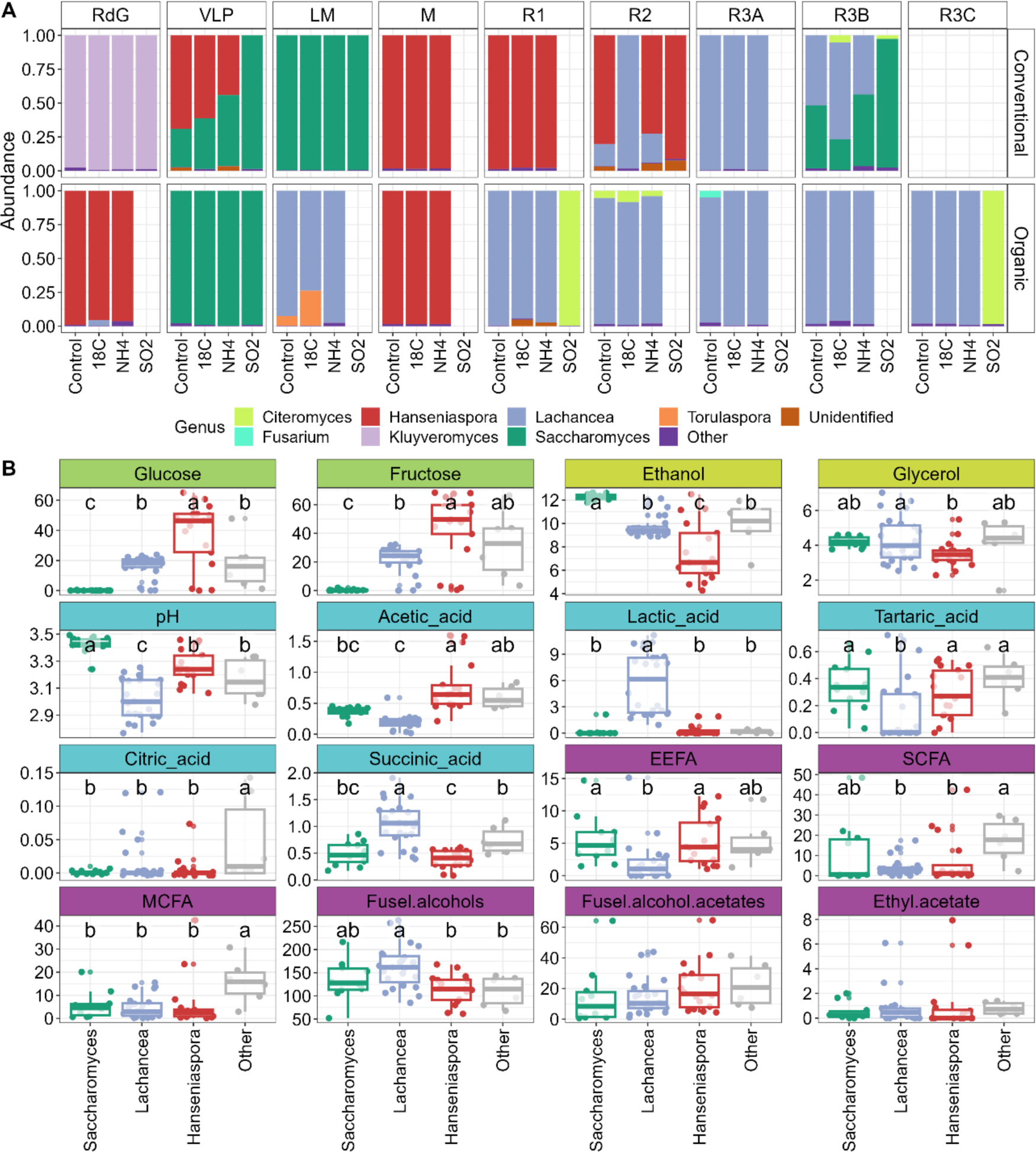
Metabolite and yeast community profiles of synthetic grape musts. **A)** Relative abundance of yeast genus during fermentations assessed via RNA sequencing. “Other” include genus with relative abundance < 2.5%. **B)** Boxplot representing the metabolite composition of fermented synthetic grape must samples. Sugars, glucose and fructose, represent the remaining concentration after fermentation, whereas the rest of metabolites are produced during this process. Vertical axis indicates metabolite concentration. Raw data detailed in **Additional File 2**. An ANOVA (Analysis of Variance) test and LSD (Least Square Difference) test were conducted (**a**– **c** indicate significance groups). *Saccharomyces*, *Lachancea*, *Hanseniaspora*, and Other presented n = 10, n = 21, n = 13, and n = 15, respectively.

On the contrary, we found distinctive metabolite profiles associated with the dominant yeast carrying the fermentation process (**Supplementary Figure S5B**, PERMANOVA: R^2^ = 0.298, *p* < 0.001). The substantial differences in the sugar consumption of fermentations dominated, and hence ethanol production, by different yeast species justify the lower concentration of aromatic compounds in fermentations dominated by non-*Saccharomyces* species (**Figure 2B**, detailed description of differences in metabolite production at different fermentation conditions can be found in **Additional File 2**). Specific differences such as the higher L-lactic acid production by *Lachancea* yeasts or the increased concentrations of acetic acid production in presence of *Hanseniaspora* have made them especially relevant in wine sciences [58–60]. When studying the molecular basis of the individual contribution of yeasts to the chemical composition of wines, most works studied individual strains in the form of pure inoculum or in co-inoculation with *S. cerevisiae*, always leading to completed fermentations [61–63]. Here, we aim to investigate the impact of different fermenting yeast populations emerged from wild complex communities, which is closer to the reality found in spontaneous wine fermentations.

### Transcriptomic profiles of fermenting yeast communities

We then evaluated the transcriptomic profiles of the fermenting communities during the most active phase (tumultuous stage) of wine fermentation. The dominant yeast species also defines the meta-transcriptome of wine fermentations (**Figure 3A**, PERMANOVA: R^2^ = 0.498, *p* < 0.001), while fermentation conditions had a much lesser impact (**Supplementary Figure S7**, PERMANOVA: R^2^ = 0.037, *p* = 0.097). Distinctive profiles emerged for samples dominated by *Saccharomyces*, *Lachancea* or *Hanseniaspora*. We used *Saccharomyces* as a control for comparative analysis, as it is the reference yeast in wine science [31]. *Hanseniaspora* presented more Differentially Expressed Orthologs (DEO) and higher accumulated Log Fold Change (LFC) than *Lachancea*, (**Figure 3BC**), 1019/7023.811 (DEO/LFC) compared to the 705/4094.062 of *Lachancea* (with 382 common DEO). *Hanseniaspora* and *Lachancea* are frequently found in grape surface and initial phases of wine fermentation, contributing to the complexity of wine aroma [64,65]. Biological processes significantly enriched in *Hanseniaspora* dominated samples were related with cell cycle and division (**Figure 3D**). This altered cell cycle could be caused by stress conditions, such as the alcohol content in the medium, which would turn lethal to *Hanseniaspora* at one point, causing the early stop of the fermentation process. *Lachancea*, on its hand, presented processes related with secondary metabolism and the production of alcohols and organic acids. These findings confirm the ability of *Lachancea* to promote the formation of higher alcohol esters, succinic acid, and reduced volatile phenols, and most importantly, L-lactic acid, as this has only been previously described for this genus [64,66].

**Figure 3.**
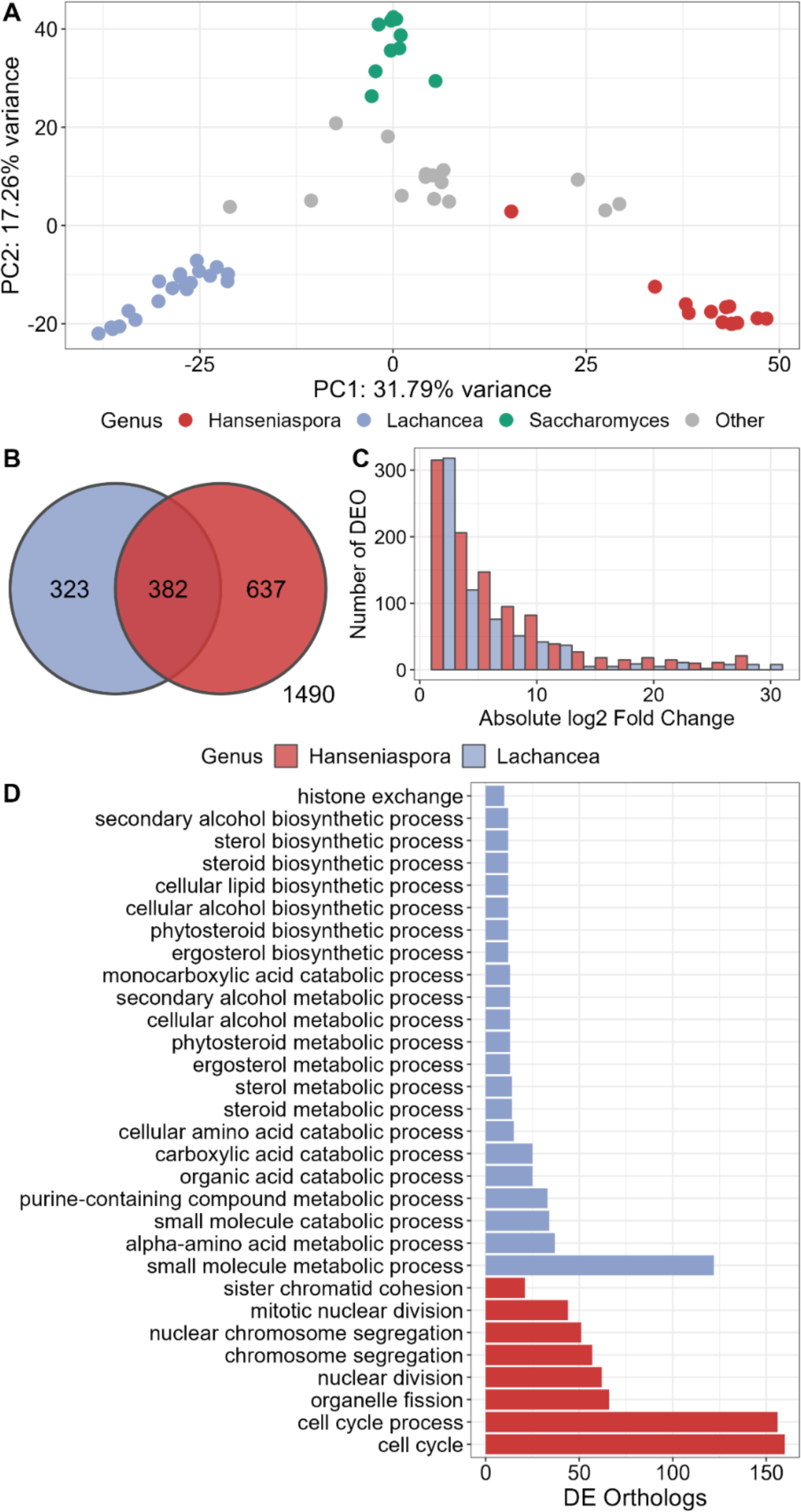
Differential expression analysis of *Hanseniaspora* and *Lachancea* dominated samples, with respect to *Saccharomyces* (*Saccharomyces*, *Lachancea*, *Hanseniaspora*, and Other presented n = 10, n = 21, n = 13, and n = 15, respectively). **A)** Principal Component Analysis (PCA) representing showing the different transcriptomic profiles colored by dominant yeast. **B)** Venn diagram comparing differently expressed orthologs (DEO) between communities dominated by *Hanseniaspora* or *Lachancea* and *Saccharomyces*. **C)** Histogram of absolute fold change (log2) expression. **D)** GO enrichment analysis of the differentially expressed orthologs.

Surprisingly, fermentative conditions had minimal influence on the meta-transcriptomic profiles, which varied significantly depending on the dominating yeasts. Only *Saccharomyces* seems to be significantly responding to the different fermentation conditions assayed. These conditions were selected because they are part of common practices in wine production [67], so it seems reasonable that winemakers have design different enological strategies that directly shape the performance of Saccharomyces, the main wine yeast. However, in a moment where spontaneous fermentations and the use of complex multi-species consortia are gaining importance in the wine industry, a deeper knowledge of the environmental preferences and metabolic traits of non-*Saccharomyces* species is needed [19].

Within the *Saccharomyces* dominated samples we observed significative effects of fermentative conditions on transcriptional profiles (**Figure 4**). Low temperature conditions caused the highest changes in transcriptomic profiles, presenting 147/329.17 DEOs and absolute accumulated log fold change, compared to 99/212.73 and 18/56.62 of NH_4_ and SO_2_, respectively (**Figure 4AB**). Low temperature mainly affected metabolic processes related to amino acids and sulfur. Cold stress would slow growth and metabolic activity, *Saccharomyces* is shown to respond by up-regulating genes involved in sulfur assimilation and glutathione biosynthesis [68]. The biological enrichment is obtained was associated with cold resistance and did not reveal any trend in metabolite production (**Figure 4C**). The addition of NH_4_ resulted in a significant modulation of the metabolism of organic nitrogen compounds and organic acids (**Supplementary Figure S8**). This translates in seemingly decreased concentrations of tartaric and succinic acids. The presence of increased concentrations of ammonia could induce nitrogen catabolite repression, inhibiting the expression of genes related with the transport of amino acids [63]. Interestingly, metabisulfite addition (SO_2_ treatments) presented little differences in transcriptomic profiles compared with control conditions, not revealing any enriched biological process. This response emerges as the result of the great tolerance of *S. cerevisiae* wine strains to metabisulfite, one of the best studied hallmarks of domestication in this species [69]. Metabisulfite addition and fermentation at lower temperatures help avoiding microbial contaminations during wine fermentation [70–73], while the addition of ammonia prevents nitrogen limitation and premature fermentation stops [74,75]. These practices are designed to ensure the safety and quality of final wines, while they seem to have a limited influence in the aroma of spontaneously fermented wines, even though they impact yeast metabolic activity.

**Figure 4.**
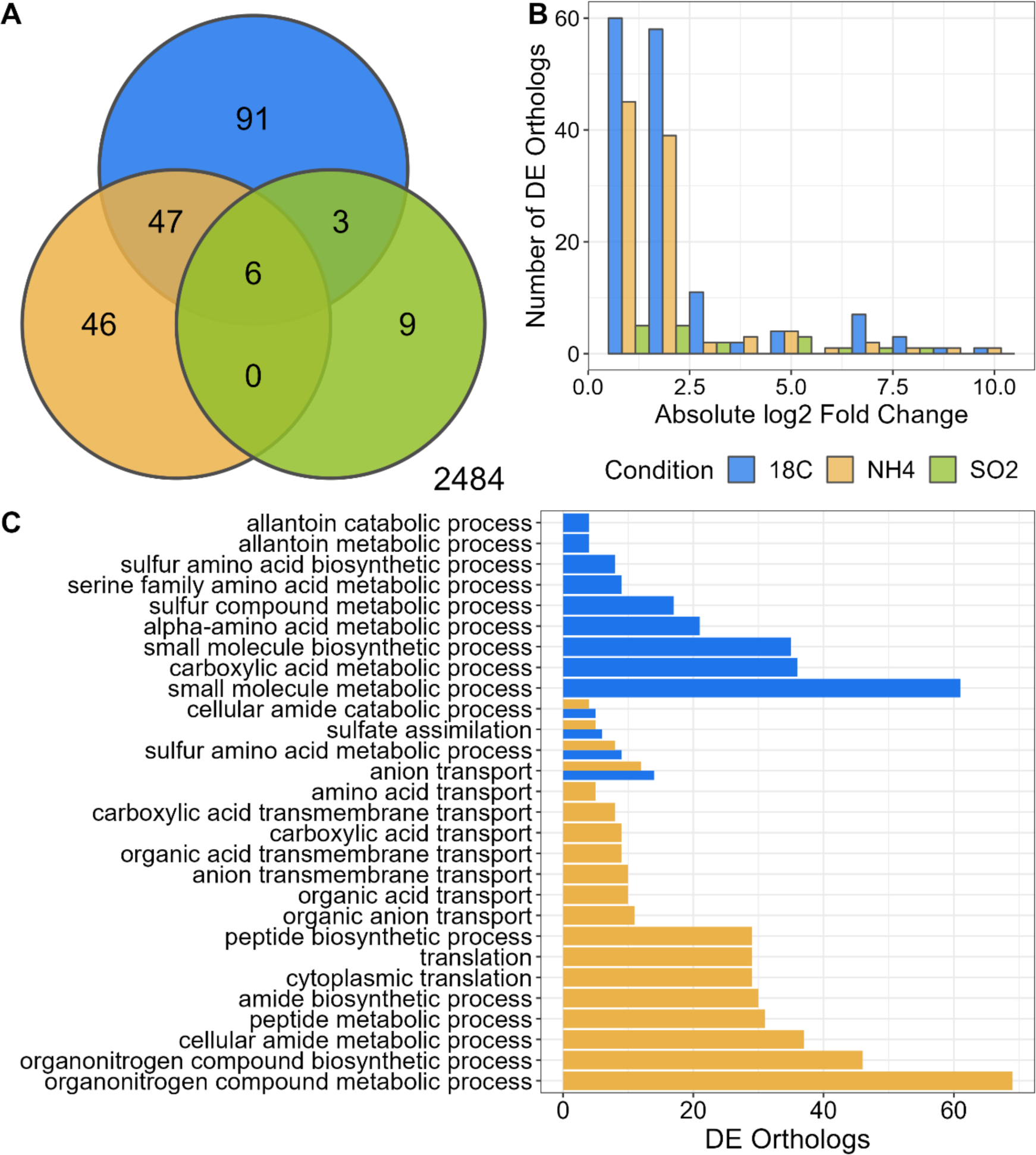
Differential expression analysis of *Saccharomyces* dominated samples comparing transcriptional profiles at different fermentative conditions (n = 2). **A)** Venn diagram comparing differently expressed (DE) orthologs across conditions. **B)** Histogram of absolute fold change (log2) expression between the experimental and control groups. **C)** GO biological process enrichment analysis of the differentially expressed orthologs. Blue represents samples fermented at low temperature (18°C), yellow samples with ammonia added (NH_4_) and green samples supplemented with metabisulfite (SO_2_).

### Orthologs responsible for metabolite production

We further aimed to understand the connection between community functional potential and metabolite production by associating the transcriptomic and metabolite profiles of the experimental fermentations. To visualize the associations between ortholog expression and metabolite production, we used bipartite networks (**Figure 5A**). Orthologs clustered into modules, each associated with specific metabolites (**Additional File 3**). Interestingly, the accumulated expression levels within each module were influenced by the dominant yeast species, reflecting distinct transcriptional profiles associated with the diverse metabolite compositions of final wines. For instance, modules 1 and 2 showed higher accumulated expression in samples dominated by *Hanseniaspora* (**Figure 5B**). These samples were characterized by higher residual sugars, acetic acid and, seemingly, fusel alcohol acetates production (**Figure 2**). The stressful fermentation conditions for *Hanseniaspora*, leading to a rapid fermentation halt, might be linked to increased ester production, such as fusel alcohol acetates [76]. In addition, *Hanseniaspora* species are also shown to produce high concentrations of acetic acid during wine fermentations [77]. The linear relationship found between the accumulated expression of module 2 and acetic acid production, even within *Hanseniaspora* dominated samples (**Figure 5B**), suggests that the expression of these orthologs is crucial for acetic acid production. Module 3 was mainly related with *Saccharomyces*-dominated samples, only correlating with ethanol content, which is highest in these samples (**Figure 2**, **Figure 5B**). Module 4 was related with *Lachancea*-dominated samples and correlated with the production of L-lactic and succinic acids, as well as fusel alcohols (**Figure 5B**). *Lachancea* outstands in the production of L-lactic and succinic acids (**Figure 2**), and the expression of these orthologs may play a crucial role on their release.

**Figure 5.**
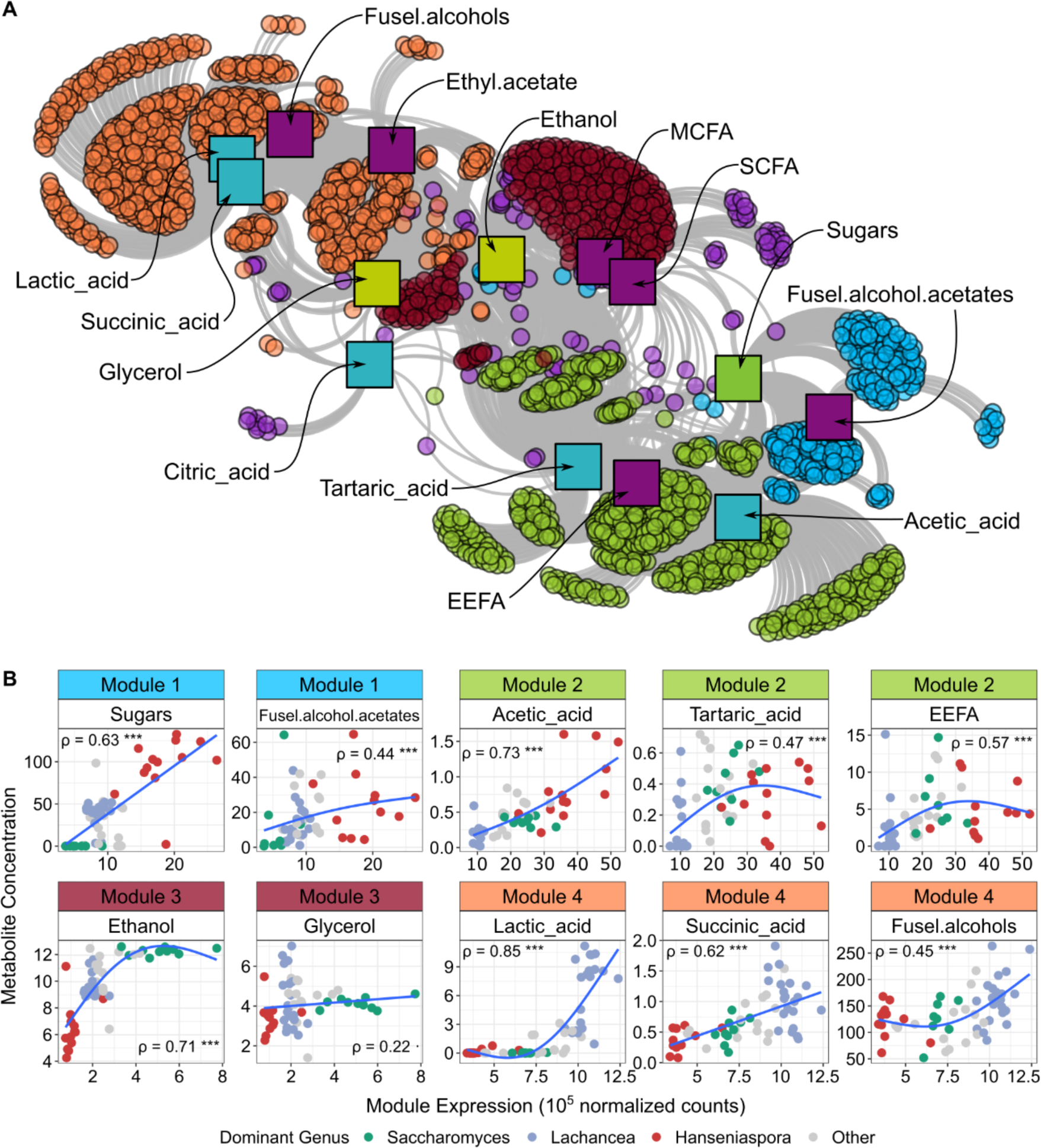
Relationship between ortholog expression and metabolite production (*Saccharomyces*, *Lachancea*, *Hanseniaspora*, and Other presented n = 10, n = 21, n = 13, and n = 15, respectively). **A)** Bipartite network showing the significant positive correlations between normalized ortholog expression and metabolite concentration. Circles represent orthologs and squares metabolites. Circle colors are indicative of module membership, i.e., orthologs associated with the same metabolites. Square colors represent the metabolite family. **B)** Relationships between the total expression of ortholog belonging to a given module, and metabolite concentration (raw data detailed in **Additional File 2**). Spearman’s rank correlation coefficient is shown (· *p* < 0.1, * *p* < 0.05, ** *p* < 0.01, *** *p* < 0.001).

We only found significant differences across fermentative conditions in the accumulated expression of module 3 (**Supplementary Figure S9**). This module is mainly expressed by *Saccharomyces*, which is prevalent in SO_2_ treated fermentations, explaining this result. The described patterns highlight differences in transcriptomic profiles among species, ultimately shaping the unique contribution of each yeast in the metabolic complexity of wine. Belda et al. [78] proposed a synthetic biology project aiming to encapsulate the functional complexity of wine communities into a single cell. Our results provide crucial insights for identifying the specific array of orthologs that define the individual contribution of yeast species to the metabolite and sensory profile of wines.

## Conclusions

In summary, our initial survey of fungal communities in fresh grape musts across diverse Spanish wine appellations, revealed significant influences of biogeography, and at a lesser extent, viticultural practices on yeast community composition. We found higher fungal diversity in vineyards under organic management. Fermenting grape must under various contrasting winemaking conditions revealed minimal effects on population dynamics and metabolite production, as the location factor differentiated initial yeast communities and must composition. The transition to an experimental setup involved the inoculation of the obtained yeast communities in synthetic grape must, revealing that variations in metabolite profiles were associated with the dominant fermenting yeast rather than fermentation conditions. Transcriptomic analyses highlighted the different profiles across yeast species, surpassing the influence of fermentation conditions. Distinctive molecular responses were observed in *Saccharomyces*, *Lachancea*, and *Hanseniaspora*, emphasizing their roles in shaping wine metabolite composition. Fermentative conditions did, however, influence the performance of *Saccharomyces*-dominated samples, which are shown to be easily conditioned by oenological standard practices. Furthermore, specific orthologs were linked to metabolite production associated with different yeast-dominated community, offering valuable insights into the functional potential of diverse yeast communities. Our findings contribute to a nuanced understanding of the intricate interplay between yeast communities, environmental conditions, and fermentation dynamics, crucial for advancing both scientific knowledge and practical applications in the wine industry.

## Declarations

### Availability of data and material

The datasets supporting the conclusions of this article are available in the NCBI Sequence Read Archive (SRA) repository: ITS PRJNA1047054 (https://www.ncbi.nlm.nih.gov/bioproject/1047054), RNA-seq PRJNA1047320 (http://www.ncbi.nlm.nih.gov/bioproject/1047320).

### Authors’ contributions

**MdC:** Conceptualization; Data curation; software; formal analysis; investigation; methodology; writing – original draft. **JR:** Conceptualization; investigation; visualization; methodology; writing – original draft. **BB-D:** Validation; investigation; methodology; writing – original draft. **JV:** Conceptualization; Investigation; methodology. **ST:** Investigation; methodology. **SI-G:** Investigation; methodology. **NR:** Investigation; methodology. **CRdV:** Investigation; methodology. **JG:** Investigation; methodology. **FZ:** Investigation; methodology. **AB:** Investigation; methodology. **LCT-C:** Investigation; methodology. **E-AL:** Investigation; methodology. **AS:** Resources; investigation. **IB:** Conceptualization; resources; supervision; funding acquisition; validation; investigation; visualization; writing – original draft; project administration. All authors read and approved the final manuscript.

## Acknowledgements

We thank the technical teams of the following wine growing companies for providing access to their vineyards and allowing the collection grape samples: Pago Los Balancines, Bodega de Las Estrellas, Bodega Dehesa de Luna, Bodega Bernabeleva, Bodega Jesús Díaz e Hijos, Bodegas Pernod Ricard.

