## Supplementary material for "Multi-omics framework to reveal the molecular determinants of fermentation performance in wine yeast populations"

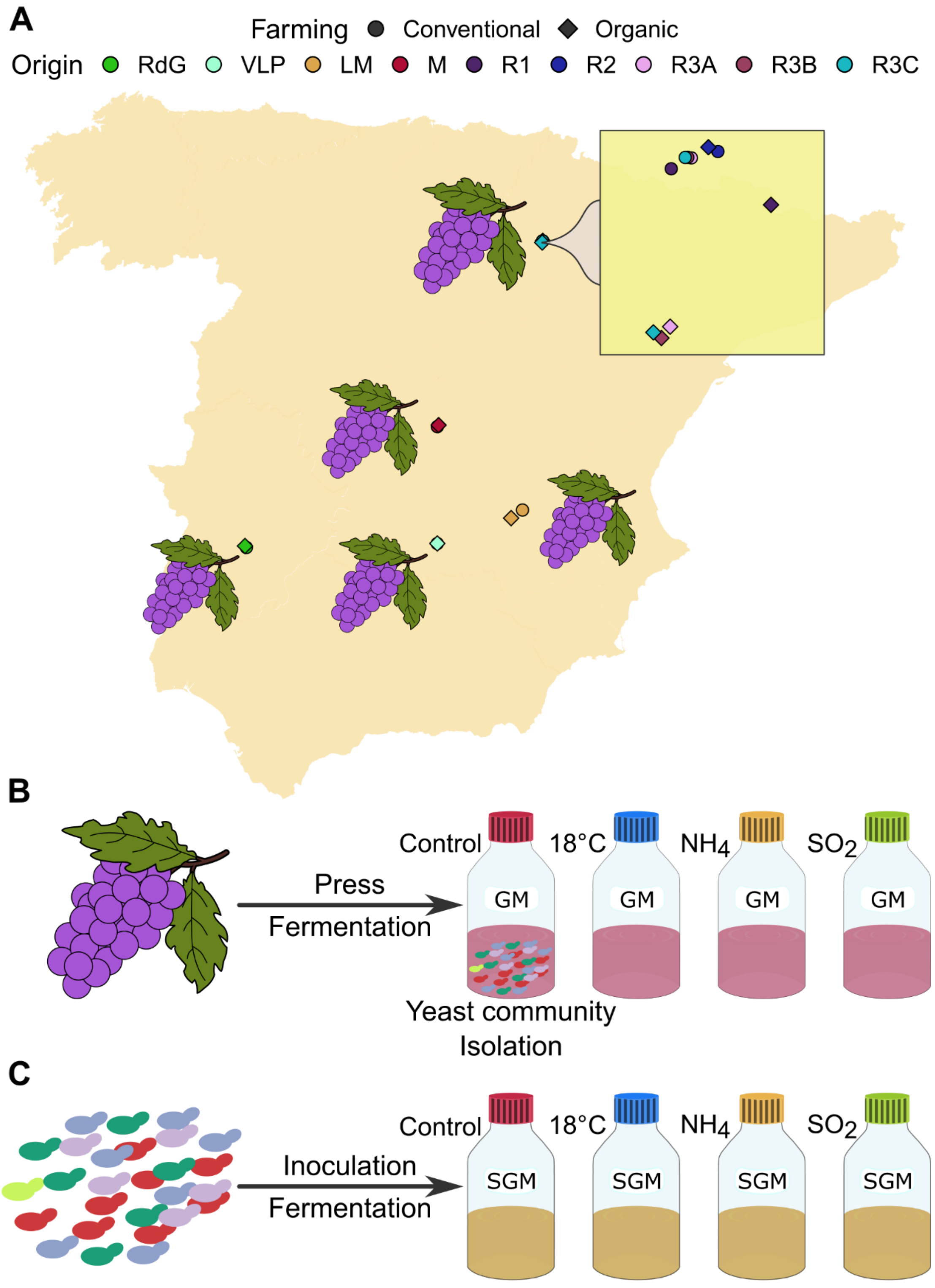

**Supplementary Figure S1.** Sampling design of the observational study and inoculation to laboratory fermentations. **A)** Sampling map representing the wine appellations (WA) surveyed and the sampling within La Rioja wine appellation. Colors represent different sampling locations: RdG (Ribera del Guadiana), VLP (Valdepeñas), LM (La Mancha), M (Madrid), R1-R3C (La Rioja, surveyed for within WA analysis). Shape indicates conventional (circle) and organic (diamond) farming managements. **B)** Observational study carried in fresh grape musts (GM). We first pressed grapes obtained at each sample point and divided the grape must in bottles, per quadruplicate. Then, each replicate was fermented under different condition until they reached the tumultuous stage, consuming between 23-45% of sugars (**Supplementary Table S1**). **C)** We repeated the fermentations in laboratory conditions by inoculating synthetic grape must (SGM) with the fermenting yeast communities obtained from control replicates of each sample.

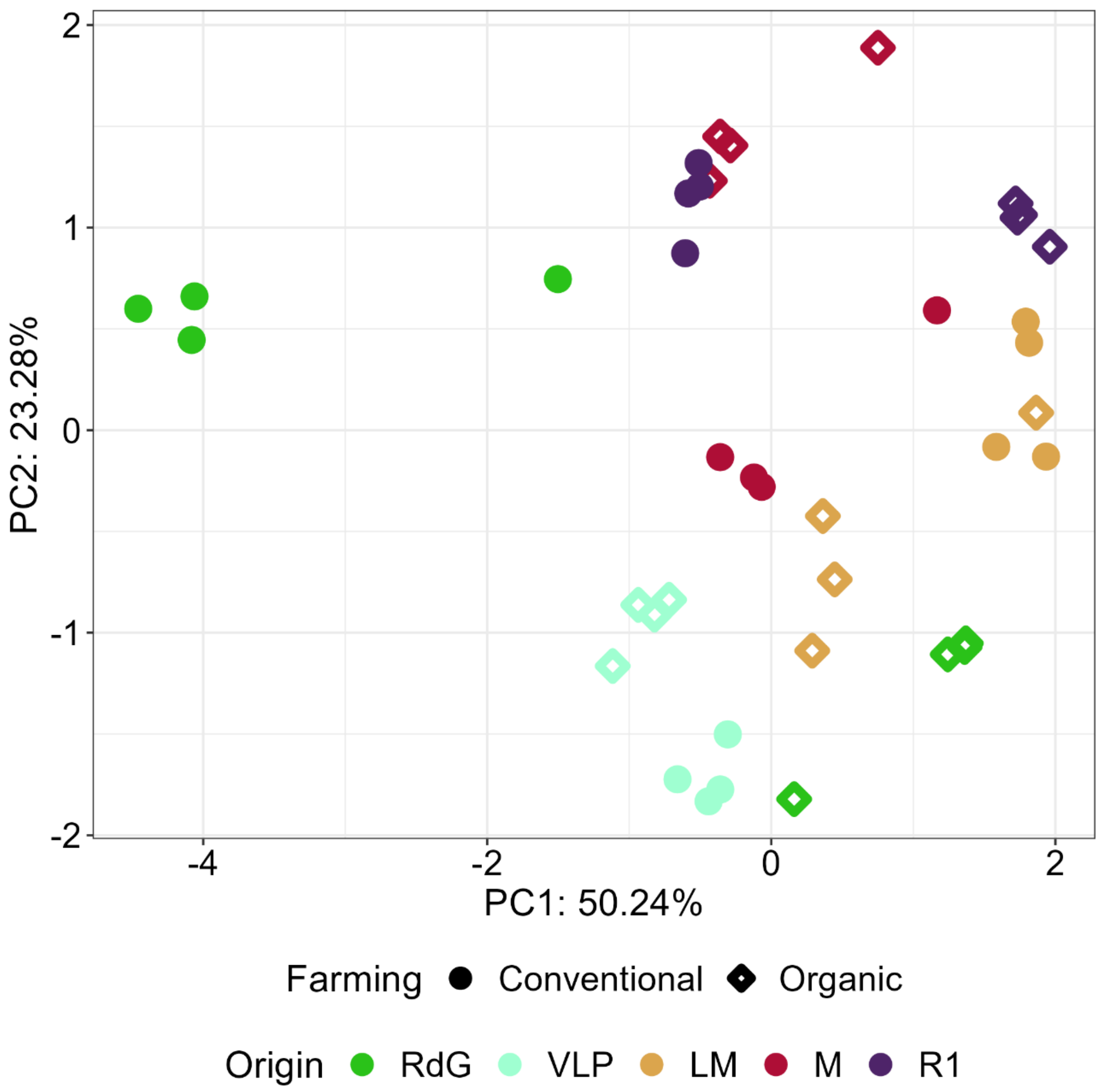


**Supplementary Figure S2.** Grape must composition of initial samples. Principal Component Analysis (PCA) representing grape must composition diversity across wine appellations, including conventional RdG samples, *i.e.,* Red Garnacha grapes (n = 4). Raw data detailed in **Additional File 2**.

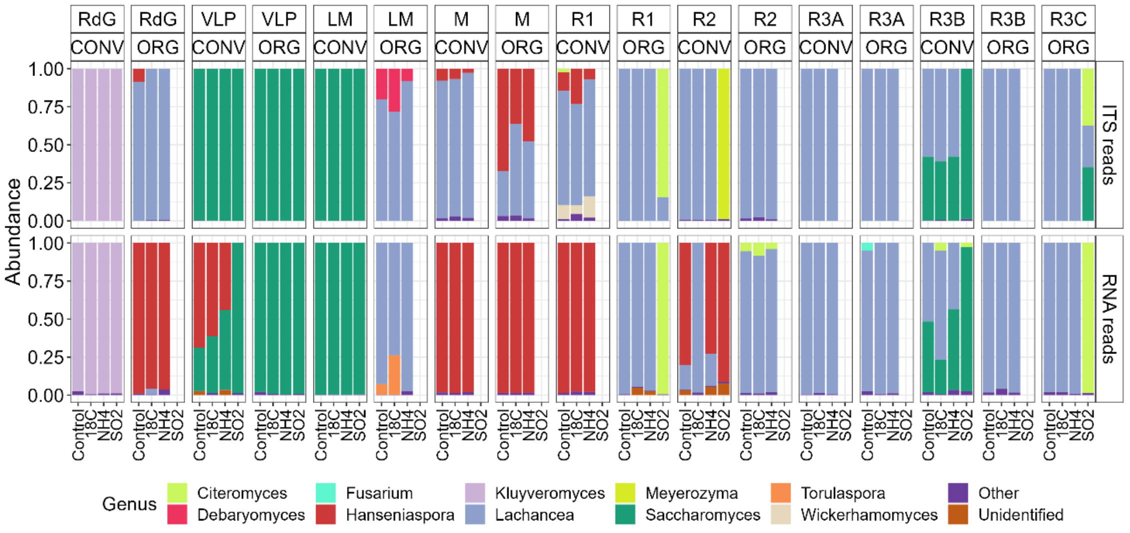


**Supplementary Figure S3.** Comparison of the taxonomic assignation achieved via ITS amplicon sequencing and meta-transcriptomics analysis. Relative abundance of fungal genus shown for each replicate from each grape sample. "Other" include genus with relative abundance <2.5%. CONV: conventional and ORG: organic farming managements.

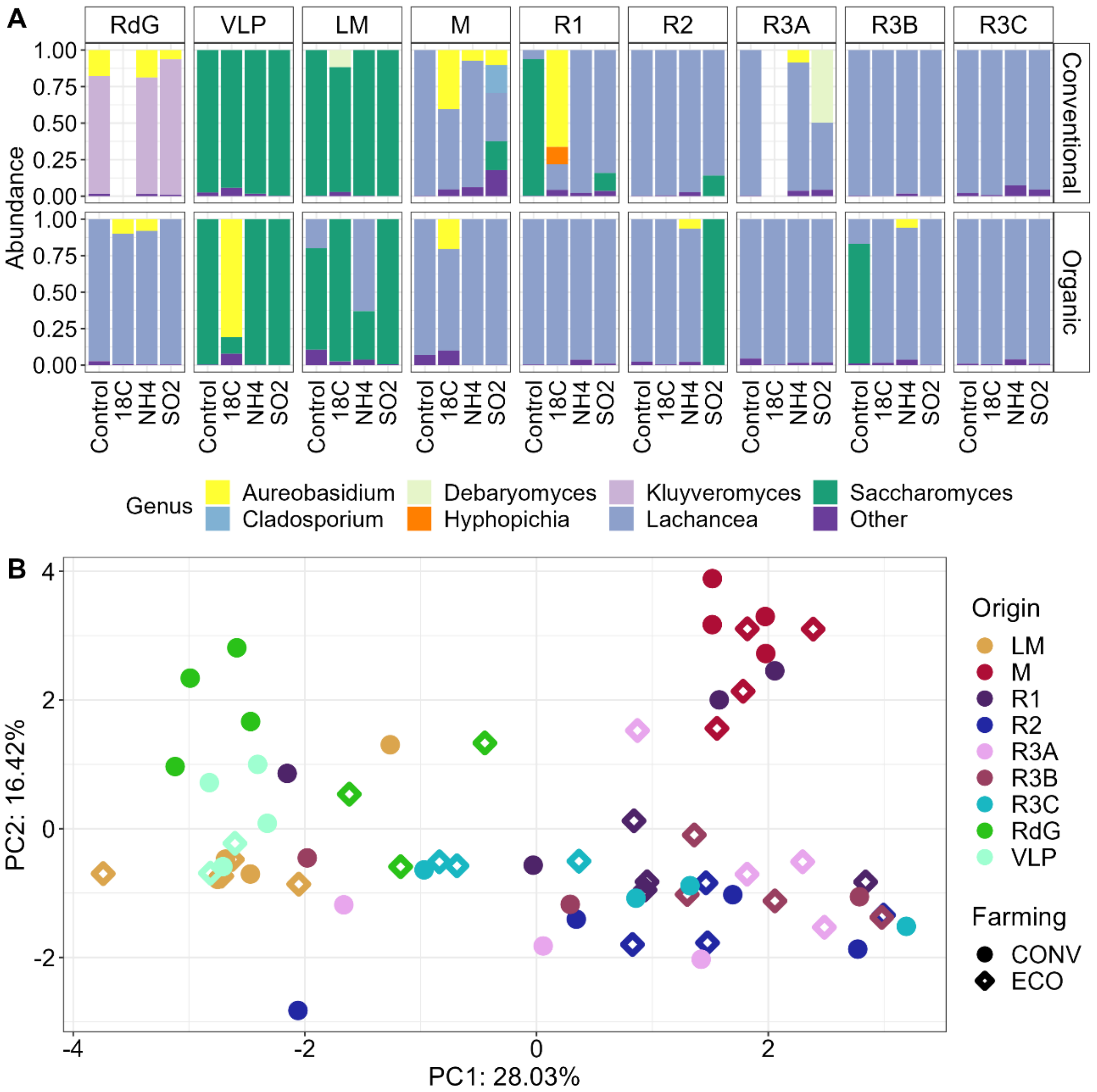


**Supplementary Figure S4.** Metabolite and yeast community profiles of fermented grape musts. **A)** Relative abundance of yeast genus during the final stage of fermentations. “Other” include genus with relative abundance < 2.5%. **B)** Metabolite profile at the end of the fermentation of fresh grape musts. Principal Component Analysis (PCA) representing metabolite profiles (n = 4). Raw data detailed in **Additional File 2**.

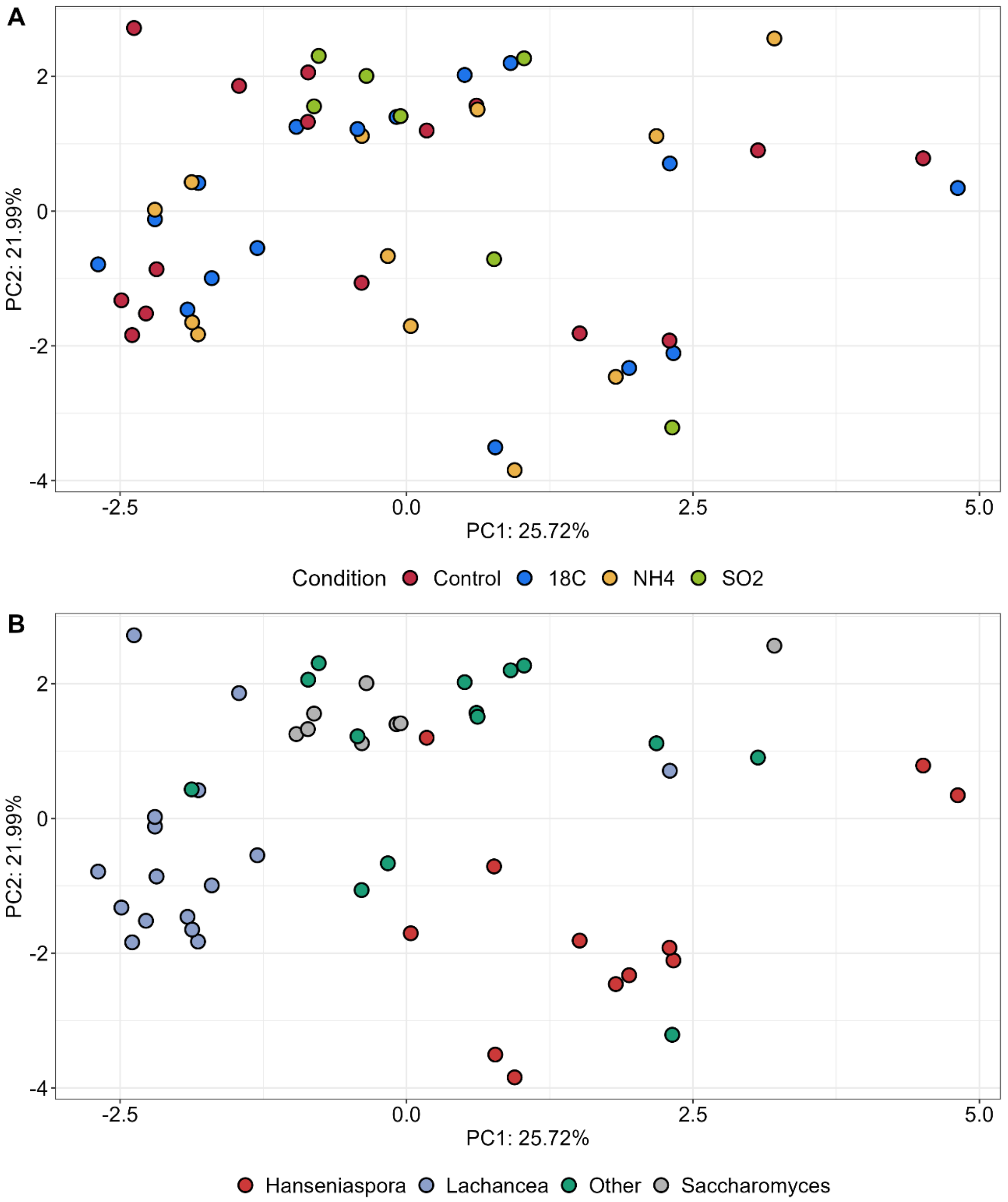


**Supplementary Figure S5.** Metabolite profile at the end of the fermentation of synthetic grape musts. Principal Component Analysis (PCA) representing metabolite profiles. Samples are colored based on **A)** fermentative conditions and (Control, 18°C, $NH_4$, and $SO_2$ conditions presented n = 17, n = 17, n = 17, and n = 8, respectively) **B)** dominant yeast (*Saccharomyces*, *Lachancea*, *Hanseniaspora*, and Other presented n = 10, n = 21, n = 13, and n = 15, respectively). Raw data detailed in **Additional File 2**.

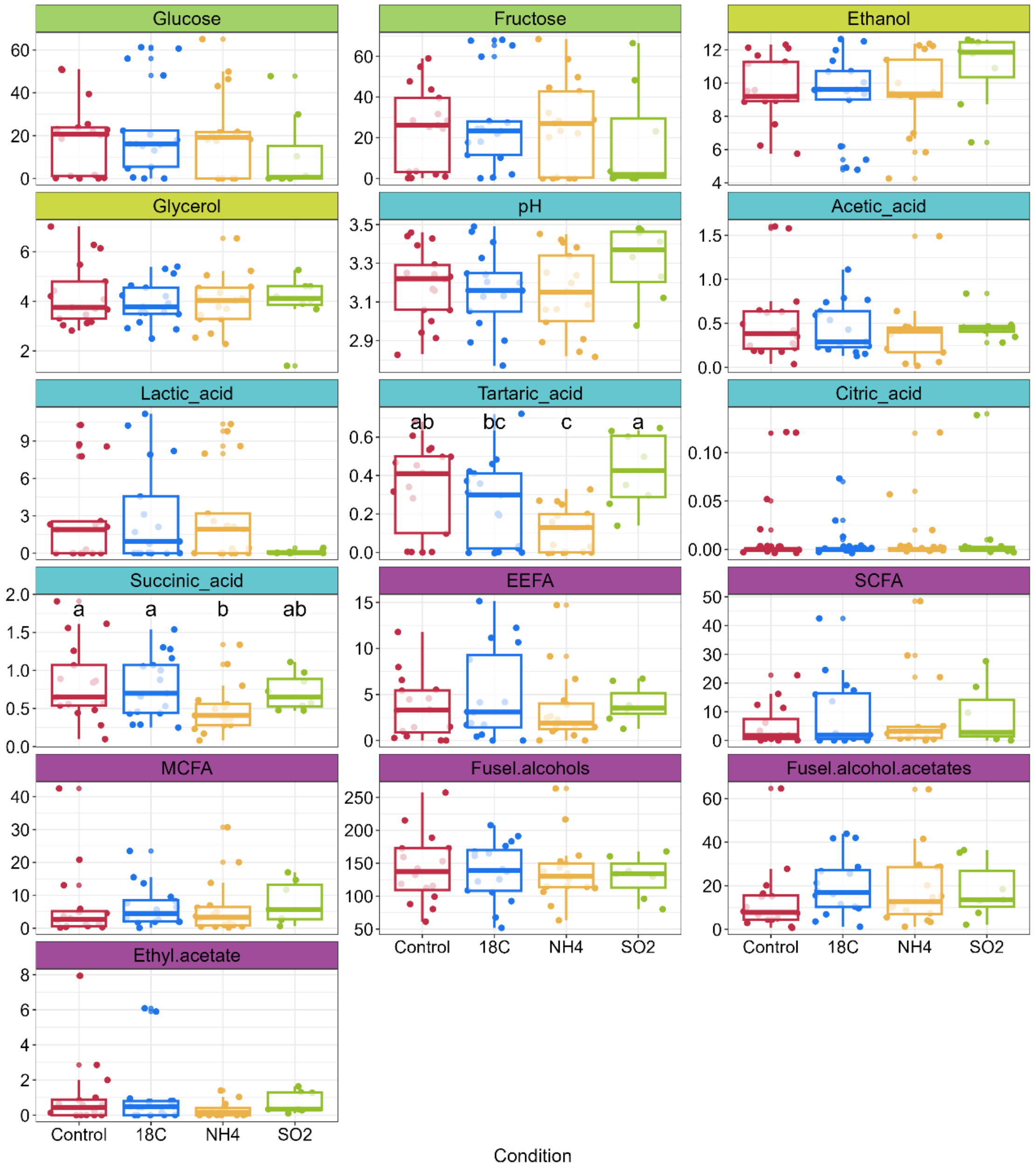


**Supplementary Figure S6.** Boxplot representing the metabolite composition of fermented synthetic grape must samples (n = 18). Sugars, glucose and fructose, represent the remaining concentration after fermentation, whereas the rest of metabolites are produced during this process. Vertical axis indicates metabolite concentration. An ANOVA test and LSD (Least Square Difference) test were conducted (**a**–**c** indicate significance groups). Raw data detailed in **Additional File 2**.

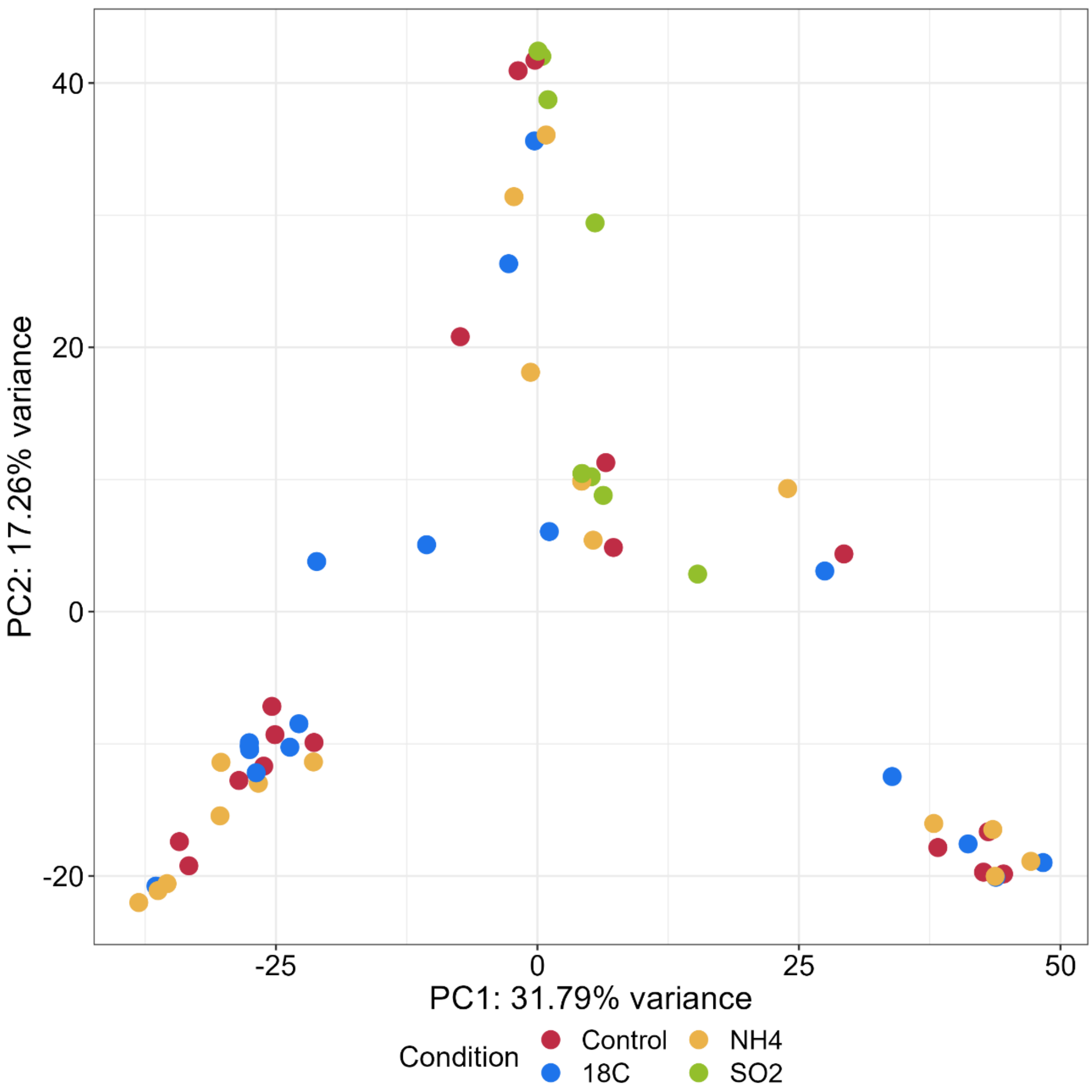


**Supplementary Figure S7.** Principal Component Analysis (PCA) showing the different transcriptomic profiles colored by fermentative conditions (Control, 18°C, $NH_4$, and $SO_2$ conditions presented n = 17, n = 17, n = 17, and n = 8, respectively).

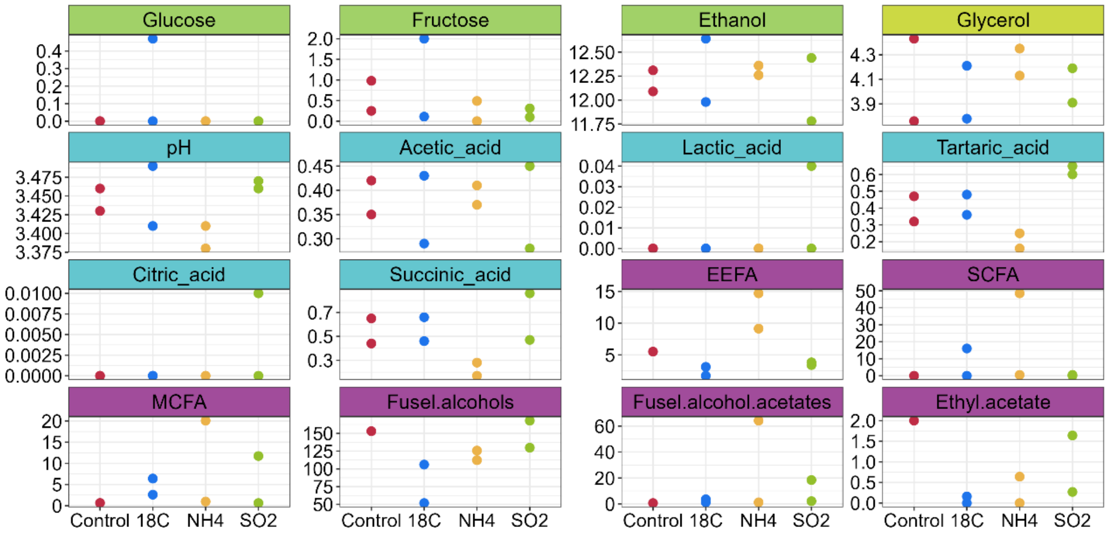


**Supplementary Figure S8.** Metabolite composition of fermented synthetic grape musts (n = 2). Glucose and fructose represent the remaining concentration after fermentation, whereas the rest of metabolites are produced during this process. Vertical axis indicates metabolite concentration. Raw data detailed in **Additional File 2**.

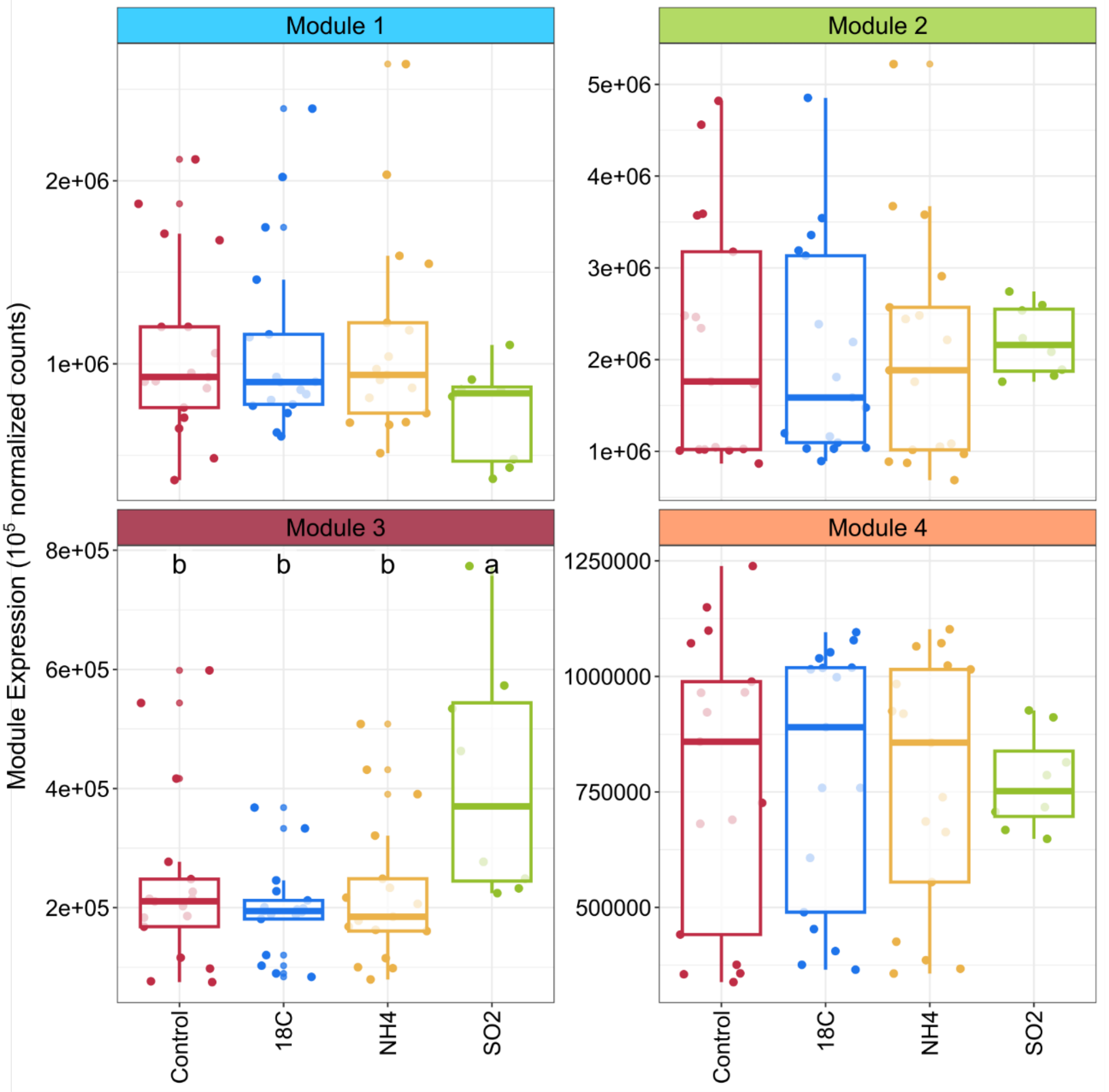


**Supplementary Figure S9**. Accumulated expression of the modules formed by orthologs in **Figure 5** (Control, 18°C, $NH_4$, and $SO_2$ conditions presented n = 17, n = 17, n = 17, and n = 8, respectively). Analysis of Variance (ANOVA) test was carried out to test for different expression levels among fermentative conditions, and further Tukey post hoc test. (a-b indicate significance levels).

**Supplementary Table S1.** Proportion of sugars consumed (%) in the sampling of yeasts communities at the tumultuous fermentation stage. Raw data detailed in **Additional File 2**.

| Sample | Grape Must | Control (SGM) | 18°C (SGM) | $NH_4$ (SGM) | $SO_2$ (SGM) |
|---|---|---|---|---|---|
| RdG-CONV | 41.67 | 19.59 | 5.56 | 11.98 | 14.28 |
| RdG-ORG | 36.52 | 17.30 | 13.79 | 17.65 | |
| VLP-CONV | 36.77 | 26.14 | 20.99 | 21.98 | 23.04 |
| VLP-ORG | 44.86 | 26.92 | 18.33 | 25.51 | 39.63 |
| LM-CONV | 32.73 | 16.43 | | 16.75 | |
| LM-ORG | 23.50 | 10.42 | 13.17 | 19.07 | |
| M-CONV | 25.22 | 34.09 | 8.96 | 17.44 | |
| M-ORG | 32.34 | 25.62 | 31.91 | 24.23 | |
| R1-CONV | 33.23 | 15.47 | 9.60 | 11.64 | |
| R1-ORG | 35.16 | 11.34 | 17.01 | 11.44 | 16.30 |
| R2-CONV | 26.12 | 24.18 | 7.95 | 17.18 | 15.46 |
| R2-ORG | 27.76 | 20.27 | 5.33 | 11.79 | |
| R3A-CONV | 33.22 | 15.96 | 8.79 | 12.93 | |
| R3A-ORG | 32.81 | 22.14 | 15.06 | 19.24 | |
| R3B-CONV | 28.57 | 31.64 | 8.82 | 20.54 | 54.44 |
| R3B-ORG | 34.84 | 20.65 | 15.83 | 14.49 | |
| R3C-CONV | 29.15 | | | | |
| R3C-ORG | 23.75 | 15.79 | 17.53 | 9.19 | 4.19 |

SGM: samples from fermentations carried in Synthetic Grape Musts. CONV: conventional; ORG: organic farming managements.

**Supplementary Table S2.** PERMANOVA results of fresh grape must composition comparing different origins and farming managements across and within wine appellations (WA).

| Scale | Source of variation | $R^2$ | F | Pr(>F) |
|---|---|---|---|---|
| Across WA | Origin | 0.547 | 23.350 | < 0.001 *** |
| Across WA | Farming | 0.043 | 70365 | 0.002 *** |
| Across WA | Origin:Farming | 0.251 | 14.304 | < 0.001 *** |
| Within WA | Origin | 0.259 | 7.410 | < 0.001 *** |
| Within WA | Farming | 0.147 | 16.811 | < 0.001 *** |
| Within WA | Origin:Farming | 0.331 | 8.470 | 0.003 ** |

**Supplementary Table S3.** PERMANOVA results comparing fungal communities comparing different origins and farming managements across and within wine appellations (WA).

| Scale | Source of variation | $R^2$ | F | Pr(>F) |
|---|---|---|---|---|
| Across WA | Origin | 0.436 | 8.160 | < 0.001 *** |
| Across WA | Farming | 0.046 | 3.412 | 0.002 ** |
| Across WA | Origin:Farming | 0.117 | 2.193 | < 0.001 *** |
| Within WA | Origin | 0.133 | 1.466 | 0.002 ** |
| Within WA | Farming | 0.070 | 3.116 | < 0.001 *** |
| Within WA | Origin:Farming | 0.119 | 1.314 | 0.018 * |

**Supplementary Table S4.** PERMANOVA results comparing fresh grape must composition comparing across different origins, farming managements across and within wine appellations and fermentation conditions.

| **Source of variation** | **$R^2$** | **F** | **Pr(>F)** |
|---|---|---|---|
| Origin | 0.483 | 10.866 | < 0.001 *** |
| Farming | 0.017 | 3.034 | 0.011 * |
| Condition | 0.055 | 3.309 | < 0.001 *** |
| Origin:Farming | 0.167 | 3.770 | < 0.001 *** |

**Additional File 1.** Raw data of metabolite composition of fresh (natural) and fermented (natural and SGM) samples.

**Additional File 2.** Supplementary discussion concerning fermented SGM metabolite composition.

**Additional File 3.** List of correlated metabolites and orthologs, indicating the ortholog module membership and KEGG annotation.
