## Additional file 2 for "Multi-omics framework to reveal the molecular determinants of fermentation performance in wine yeast populations"

**Additional File 1**

**Lower concentration of tartaric and succinic acids in presence of ammonia**

We did not observe a conserved pattern in metabolite production during experimental fermentations under contrasting conditions: Control, 18°C, NH_4_ and SO_2_ (**Supplementary Figure S5A**, PERMANOVA: R^2^ = 0.062, *p* = 0.421). We only found a general decrease in the final concentration of tartaric and succinic acids in the NH_4_ condition, adding diammonium sulphate as a nitrogen nutrient (**Supplementary Figure S6**). Tartaric acid comes from grape musts and microbes are unable to metabolize it, but could precipitate as less soluble ammonium tartrate salts [1–4]. Succinic acid is produced as a by-product of yeast metabolism during fermentation, and it has been reported to be dependent of the nitrogen source, with the addition of ammonia yielding the worst results [5,6].

**Dominant yeasts produce contrasting metabolite profiles during experimental fermentations**

On the contrary, we found distinctive metabolite profiles associated with the dominant yeast carrying the fermentation process (**Supplementary Figure S5B**, PERMANOVA: R^2^ = 0.298, *p* < 0.001). Since we performed our trials without the typical contamination of winery facilities, where *Saccharomyces* is often inoculated in spontaneous fermentations, many samples were dominated by whether *Lachancea* or *Hanseniaspora*. However, due to their lower ethanol tolerance compared with *Saccharomyces* [7–9], they were not able to end fermentations, resulting in lower sugar consumption and ethanol production (**Figure 2B**). *Lachancea* and *Hanseniaspora* showed higher yields of glycerol (**Figure 2B**), which is a by-product of wine fermentation [10]. The wine industry values higher glycerol production because it can result in wines with lower ethanol contents and improved mouthfeel attributes [11], making *Lachancea* and *Hanseniaspora* strains with high glycerol yields attractive for wine production [12–14]. *Lachancea* produced higher acidity, which is an important factor defining wine quality [15], because of higher L-lactic and succinic acids production, whereas *Hanseniaspora* produced more acetic acid (**Figure 2B**). The inoculation of *Lachancea* strains is proposed because of its ability to produce L-lactic and succinic acids, a trait that gains importance as a strategy to mitigate the impact of climate change in wine acidity [16]. Volatile compounds play important roles in shaping wine aroma [17], and its production was overall higher in *Saccharomyces* dominated fermentations, highlighting the also relatively high production by other minoritarian yeasts, like *Kluyveromyces*. The metabolic activities of native non-*Saccharomyces* yeasts are expected to increase the aromatic complexity of wines when used as mixed inoculum with *S. cerevisiae* [18]. The substantial differences in the sugar consumption of fermentations dominated by different yeast species justify the lower concentration of aromatic compounds in fermentations dominated by non-*Saccharomyces* species. Most studies on the contribution of yeasts to the chemical composition of wines are focused on the use of single species. Here, we investigate the impact of different yeast species within a context of complex communities of various yeast species, which is closer to the reality found in the wine fermentation industry.
